# HiC-inspector: a toolkit for high-throughput chromosome capture data

**DOI:** 10.1101/020636

**Authors:** Giancarlo Castellano, François Le Dily, Antonio Hermoso Pulido, Miguel Beato, Guglielmo Roma

## Abstract

HiC-inspector is a toolkit for the analysis and visualization of data generated by high-throughput chromatin conformation capture (HiC). The analysis module comprises steps of data formatting, genome alignment, quality control and filtering, identification of genome-wide chromatin interactions, and statistics. The interactive browser enables visual inspection of interaction data generated and analysis results.

## INTRODUCTION

It has long been known that transcription regulation is related to chromosomal organization and to the 3-dimensional architecture of the genome. Among available methods, high-throughput conformation capture (HiC) allows to study the three-dimensional architecture of whole genomes through the detection of long-range chromosomal interactions [PMID: 20461051]. This technique consists of cross-linking of cells with formaldehyde resulting in covalent links between spatially adjacent chromatin segments, digestion of chromatin with a restriction enzyme, ligation of the resulting ends, and tagging of interacting fragments by paired-end sequencing. Sequencing reads generated can be therefore analyzed for the identification of genome-wide interactions; however, like for other high-throughput technologies, the stage of data processing and analysis (rather than data production) remains the major bottleneck for the researchers. Here, we present HiC-inspector, a toolkit to facilitate the analysis and the visualization of HiC data. The analysis pipeline performs read alignment, filtering of paired-end reads that are in the DNA fragment size window around the restriction enzyme sites, counting of interactions with a user-defined resolution –which allows filtering of artifactual fragment pairs-, and generation of contact matrices and heatmaps. As a result, an interactive browser is offered to help visualize the mapping of the observed expected and corrected interactions in the reference genome. The software is open source and available at: http://biocore.crg.cat/wiki/HiC-inspector.

## METHODS

The input for the HiC-inspector can be the standard sequence output files of any massively parallel DNA sequencing platform, even in compressed format. Two files of paired-end sequencing data have to be provided, one for each mate of sequences. The format of the input files is kept quite flexible. As first option, reads provided in qseq or fastq format are aligned to the genome of interest using the program Bowtie (Langmead B, et al. 2009), allowing only ungapped matches and no more than two mismatches. A statistics of the mapping is provided in the output. As alternative, bed files corresponding to the genomic positions of the paired-end sequencing reads previously aligned to the reference genome can be provided, which makes HiC-inspector compatible with all of sequencing platforms and alignment tools. In both formats, raw or aligned, a unique name for each sequence mate is required.

As first quality criterion, HiC-inspector ensures that aligned reads are the results of proximity-based ligation of digested fragments. To this purpose, both reads in a pair must align to the genome near the restriction sites, and, in particular, within a distance lower than the size of the DNA fragment after sonication. Furthermore, only reads with correct orientation toward the restriction sites are kept (3’-end of both reads facing the restriction site). These quality control steps are performed with BEDTools utilities (Quinlan AR et al., 2010) that are built in our program. A percentage of intra- and inter-chromosomal interactions identified and the frequency distribution of paired reads distances are provided in the output.

## RESULTS

Only reads passing the quality control are considered in the generation of contact matrices of chromatin interactions. During the analysis, users can define the desired genomic resolution, e.g. the genomic windows, or bins (default 1Mb), to split the genome and count for number of interactions. Several genomic bins can be provided as input, thus allowing the generation of more contact matrices in the same analysis. A visualization of these matrices is provided with colored heat-maps, where color intensities correlate with the frequencies of interactions and revealing which part of the chromosomes are “close to” or “far away from” each other. As final results, matrices of observed interactions, matrices of interactions corrected for genome coverage and Pearson correlation matrices are generated.

The correction for coverage is calculated according to the following formula:

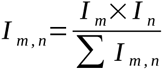

where I is the number of observed interactions and m and n indicate the bin coordinates in the matrix. For intra-chromosomal interactions, a matrix of expected interactions is generated, where the expected number of interactions is determined by the average contact probability:

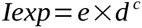

where Iexp represents the expected number of interactions and d the genomic distance between the paired reads. The coefficients e and c are derived by the frequency distribution of the paired read distances. The observed /expected matrix is then obtained. The Pearson correlation matrices are generated by calculating the Pearson correlation coefficient in each bin between the horizontal and vertical values of the corresponding bin.

All possible matrices and heat-maps are produced for the whole genome as well as for each single chromosome and chromosome pairs. As final results, HiC-inspector provides summary statistics concerning the frequency distribution of paired reads distances, that is necessary to estimates the expected interactions, and a bar plot for quality check containing the number of reads in the total dataset, aligned to the genome, filtered after QC and belonging to intra- or inter-chromosomal interactions.

### Interactive browser

The analysis pipeline generates a simple interactive HTML report to provide a visual representation of the results. All different chromosome contacts combinations can be browsed as image graphs next to each other within the same page. The relative heat maps of color intensity are obtained using the “heatmap.2” function from the {gplots} library in R (http://www.r-project.org/) and represent the contact frequency for each segment of the genome. Likewise, researchers can easily use it for sharing their results out of the box online with the scientific community.

### Implementation

HiC-inspector has been compiled in a way that it can be easily extended with new algorithms developed for the Hi-C applications. To test and illustrate its performance, we tried our toolkit on a 8 CPU (4000 Bogomips each) and 48 GB RAM workstation and processed the data derived from Lieberman A. et al. downloaded from the Sequence Read Archive (Accessions: SRX011608, SRX011609, SRX011610, SRX011611). Final results, starting from FASTQ files on Hg19 genome, took an average of 2 ¼ hours. The heat maps obtained with HiC-inspector are reproducible according to what is depicted in the Hi-C Data Browser (hg18) available at http://hic.umassmed.edu.

HiC-inspector can moreover be implemented with the recently published bioconductor package HiTC by Servant N et al. (2012). This allow to combine a very a fast way of processing the raw data by HiC-inspector and after transformation, normalization and annotation with the HiTC package the resulting interaction maps can be easily visualized in the interactive HiC-inspector browser.

## CONCLUSIONS

HiC-inspector is an open-source software tool to analyze high-throughput conformation capture (Hi-C) data and investigate the three-dimensional architecture of whole genomes. Our computational pipeline is a useful solution for automated high-throughput computational support of Hi-C genome sequencing projects. We aim to provide a user-friendly tool able to perform full-automated and highly complex computational HiC analyses, as well as an interactive browser to proper visualize the resulting long-range chromosomal interactions. This can be used to generate pictures for publications, which can be easily shared with collaborators as HTML reports. The modular organization of HiC-inspector allows integration of new developed algorithms, thus making it possible to stay up-to-date with developments in high-throughput Chromosome Conformation Capture.

## ACKNOWLEDGEMENTS

The authors thank the Chromatin and Gene Expression lab and the Bioinformatics Core Facility at the Centre for Genomic Regulation (CRG) for the useful discussions.

---

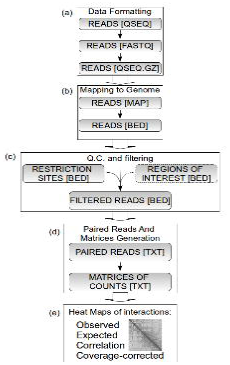

